# Signaling from the *C. elegans* Hypodermis Non-autonomously Facilitates Short-term Associative Memory

**DOI:** 10.1101/2023.02.16.528821

**Authors:** Shiyi Zhou, Yanping Zhang, Rachel Kaletsky, Erik Toraason, Wenhong Zhang, Meng-Qiu Dong, Coleen T. Murphy

## Abstract

Memory loss is one of the most debilitating symptoms of aging. While we normally think of memory regulation as autonomous to the brain, other factors outside of the brain can also affect neuron function. We have shown that the longevity insulin/IGF-1 signaling (IIS) pathway regulates neuron-specific activity, but whether the maintenance of memory through the IIS pathway is completely autonomous to the nervous system, or if there are systemic inputs, is unknown. We address this question using the auxin-inducible degradation (AID) system to degrade the IIS receptor, DAF-2, in specific tissues. Surprisingly, DAF-2 degradation specifically in the hypodermis improves memory in both young and aged *C. elegans* through the hypodermal expression of the diffusible Notch ligand, OSM-11. The Notch receptor LIN-12 is required for OSM-11’s effect on memory, as are the downstream transcription factor LAG-1 and co-activator SEL-8. Furthermore, mid-life overexpression of the Notch ligand OSM-11 improves memory and slows cognitive decline. Hypodermal DAF-2 degradation suppresses the expression of *ins-19*, whose downregulation extends memory. Together, our data suggest a model in which the hypodermis, a metabolic tissue, can non-autonomously regulate neuronal activity and function, indicating a systemic connection between metabolism and memory regulation.

## Introduction

As human lifespan continues to increase^1^, an emerging challenge must be addressed: how can we slow cognitive aging? Age-related cognitive decline starts from middle age^2^, with learning and memory processes decreasing more rapidly than other behavioral and motor declines, even in the absence of disease^3,4^. While mechanisms underlying memory regulation and cognitive aging have been studied for decades, therapeutic treatments for age-related cognitive decline are still lacking.

*C. elegans* is a useful model for studying cognitive decline because the molecular machinery required to form and maintain short and long-term memories is conserved^5,6^. For example, pathways controlling presynaptic vesicle transport and release are key components regulating memory maintenance in both worms and humans^7–9^. Like mammals, *C. elegans* can learn and form short- and long-term associative memory^3,6^. Short-term memory in worms can be measured using a Pavlovian-style appetitive associative learning and memory assay (STAM)^3^. Briefly, worms’ preference for an odorant (butanone) is tested at selected time points after conditioning training in the presence of both food and butanone to measure short-term associative memory. We previously found that young adult worms retain this short-term memory for about two hours. This memory capacity noticeably decreases with age and is ultimately lost by midlife (Day 6) of adulthood^3^.

The insulin/IGF-1 signaling (IIS) longevity pathway has been found to be critical in regulating cognitive aging in worms and mammals^7^. This pathway both autonomously and non-autonomously regulates longevity, metabolism, and reproduction^10–13^. In addition, the reduction of IIS signaling via mutagenesis of the IIS receptor, *daf-2*, slows cognitive aging in worms^3^. While genes responsible for neuron-autonomous regulation of this memory improvement have been identified by transcriptomic analysis of neurons isolated from *daf-2* mutants^8^, it is unclear whether there is non-autonomous short-term memory regulation by other tissues, and whether such regulation is mechanistically distinct from that of direct neuronal inputs.

To investigate whether there are inputs from non-neuronal tissues to enhance short-term memory, we used the auxin-induced protein degradation system (AID) to selectively degrade DAF-2 in the major tissues of the worm^14^ and we subsequently tested these animals for their short-term associative memory (STAM). In addition to the expected memory improvement by neuronal DAF-2 degradation, we were surprised to find that hypodermal DAF-2 degradation significantly improves short-term memory. We further established that this memory regulation by the hypodermis is mediated by Notch ligand signaling from the hypodermis to neurons, and subsequent gene transcription changes lead to improved memory. Furthermore, the reduction of hypodermal IIS or increased Notch signaling can each rescue short-term memory decline in aged worms. Together, our data establish a role for non-autonomous regulation of memory and cognitive decline by hypodermal IIS signaling and downstream neuronal Notch signaling to neurons.

## Results

### Reduced Insulin Signaling in the Hypodermis Extends Short-term Memory

We previously found that reduction of insulin/IGF-1 signaling improves *C. elegans* short-term associative memory in both young and aged adults^3^. Consistent with our previous results, the insulin-like receptor mutant *daf-2* improved memory maintenance in young Day 1 adult worms, retaining the memory beyond two hours post-conditioning, while wild-type worms had forgotten the association between food and butanone^3^ by 90 minutes (Supplementary Figure S1A). To determine whether this enhanced memory maintenance was due solely to *daf-2*’s tissue autonomous effects within the nervous system or if there are systemic inputs from reduced IIS in other tissues, we used the auxin-induced protein degradation system (AID) to selectively degrade DAF-2 in individual tissues with temporal control. This system allows us to degrade DAF-2 only in specific tissues in adult animals^14^, avoiding any possible developmental effects due to reduced IIS. Using auxin treatment of Day 1 adult worms, followed by the Pavlovian appetitive short-term associative memory (STAM) assay on Day 2 (young adults) (Supplementary Figure S1B), we examined short-term associative memory after DAF-2 was degraded in neurons, intestine, gonadal sheath, germline, muscle, or hypodermis (**Figure 1**). We found that neuronal DAF-2 degradation significantly improved memory duration (**Figure 1A**), consistent with the role of neuron-autonomous IIS on memory and behavior^8^. Degradation of DAF-2 in the intestine, gonadal sheath, or germline had no effect on short-term associative memory (**Figure 1B-D**). The muscle DAF-2 AID strain shows significant defects in learning prior to AID treatment (Supplementary Figure S1C), making it challenging to test short-term memory and complicating interpretation. Surprisingly, hypodermal DAF-2 degradation dramatically enhanced memory duration (**Figure 1E**). The magnitude of memory improvement induced by hypodermal DAF-2 degradation was similar to that observed with neuronal DAF-2 degradation (**Figure 1F**).

**Figure 1.**
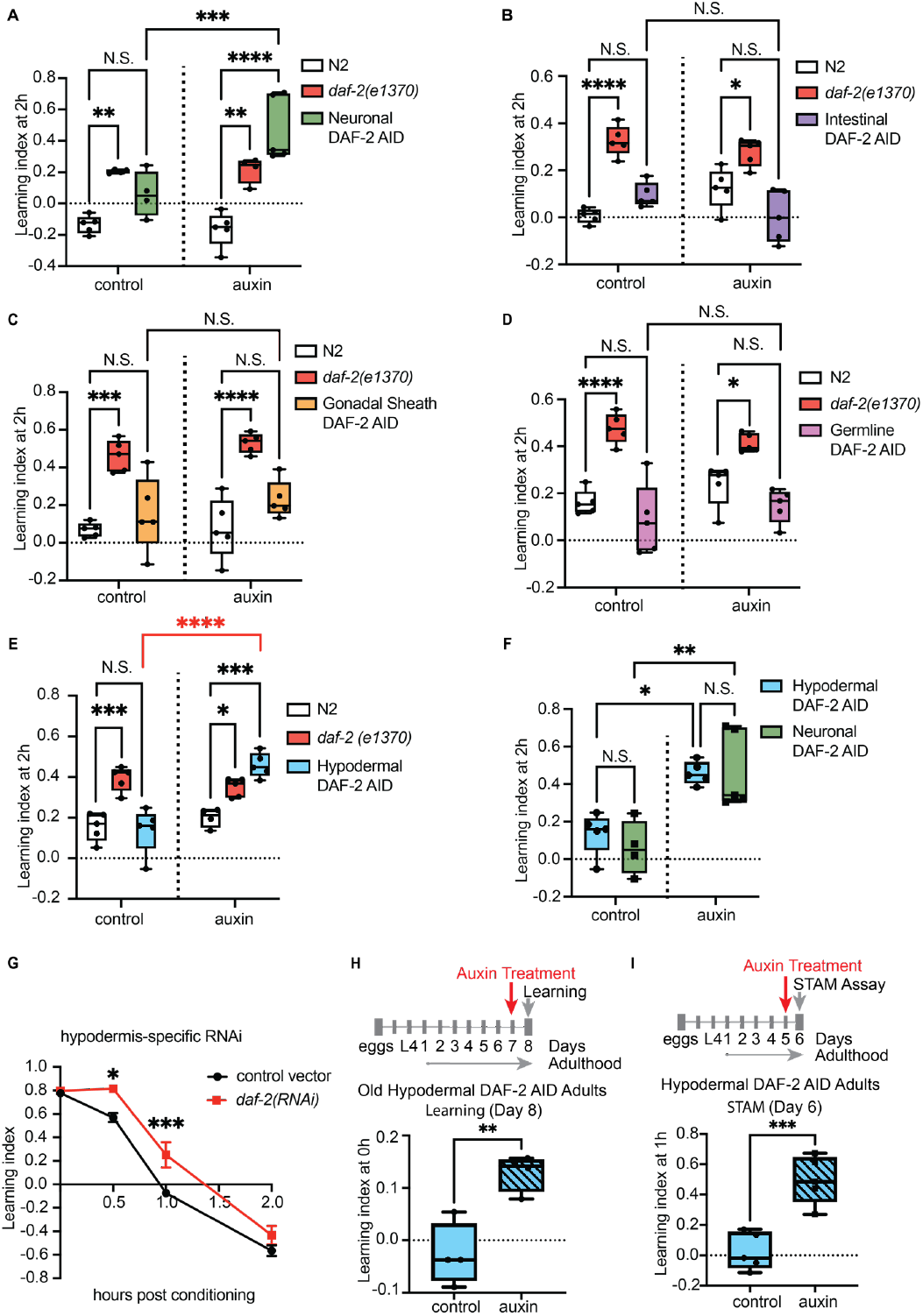
DAF-2 Degradation in the Hypodermis Improves Memory in Young and Aged Worms. (A) Auxin-induced DAF-2 degradation specifically in neurons increased short-term associative memory in Day 2 adult animals (MQD2356). Auxin treatment was started on Day 1 for all experiments unless indicated otherwise. Representative experiment for 2 biological replicates. Two-way repeated measures ANOVA, Tukey post hoc tests. (B-D) Auxin-induced DAF-2 degradation in the other tissues (intestine: MQD2374; gonadal sheath: MQD2383; germline: MQD2375) did not affect short-term associative memory. Two-way repeated measures ANOVA, Tukey post hoc tests. (E) Auxin-induced DAF-2 degradation specifically in the hypodermis improved short-term associative memory in Day 2 adult animals (MQD2378). Representative experiment for 3 biological replicates. Two-way repeated measures ANOVA, Tukey post hoc tests. (F) Comparison between memory improvements induced by hypodermal and neuronal DAF-2 degradation. Two-way repeated measures ANOVA, Tukey post hoc tests. (G) Knocking down *daf-2* in young adult hypodermal-specific-RNAi worms (IG1846) increased short-term associative memory. Representative experiment for 3 biological replicates. Mean ± s.e.m., two-way repeated measures ANOVA, Bonferroni post hoc tests. (H) Auxin treatment was started on Day 7, and STAM assay was performed on Day 8. Auxin-induced DAF-2 degradation, specifically in the hypodermis (MQD2378), improved learning in Day 8 adult animals. Representative experiment for 3 biological replicates. Unpaired t-test. (I) Auxin treatment was started on Day 5, and STAM assay was performed on Day 6. Auxin-induced DAF-2 degradation, specifically in the hypodermis (MQD2378), improved memory at 1hr post-conditioning in Day 6 adult animals. Representative experiment for 3 biological replicates. Unpaired t-test. In all experiments, n ≥ 4. (n represents the number of chemotaxis plates at each time point. Each plate contains ∼100 worms.) *p ≤ 0.05, **p ≤ 0.01, ***p ≤ 0.001, ****p ≤ 0.0001.

To confirm that auxin-induced reduction of DAF-2 was specific to the hypodermis, we examined DAF-2::degron::mNeonGreen fluorescence after auxin treatment and confirmed that DAF-2 protein was specifically degraded only in the hypodermis and not in neurons (Supplementary Figure S1D). To further test the role of hypodermal DAF-2 in short-term associative memory, we knocked down *daf-2* in a strain that is competent for RNA interference (RNAi) only in hypodermal cells; hypodermis-only knockdown of *daf-2* was sufficient to increase memory performance following training (**Figure 1G**), confirming a role for hypodermal IIS in memory regulation in young worms.

We previously found that aged worms lose the ability to learn and to remember with age, but reduction of insulin signaling results in prolonged maintenance of learning and short-term associative memory^3^. However, these experiments were performed using mutants in which IIS was reduced in all tissues^3,8^. It is unclear whether the reduction of DAF-2 in specific tissues later in the worm’s life, particularly when animals are already experiencing cognitive decline, can preserve or ameliorate aging-induced memory loss. To address this question, we degraded DAF-2 in the hypodermis in mid-life, when cognition has already declined, and then tested the worms’ learning and memory. We found that acute DAF-2 degradation in the hypodermis improved learning in aged worms (Day 8) (**Figure 1H**). To test memory, which declines earlier with age than learning^3^, we started auxin treatment on Day 5 and performed memory assays on Day 6, when *daf-2* mutants are still capable of learning and short-term memory, but wild-type worms are not. Mid-life (Day 5) DAF-2 degradation specifically in the hypodermis significantly increased short-term associative memory in Day 6 worms (**Figure 1I**), indicating beneficial effects from the reduction of hypodermal IIS in cognitive aging. Together, these results suggest that reducing hypodermal insulin signaling via DAF-2 degradation specifically in the hypodermis enhances short-term associative memory in both young and aged adults.

### Identification of Hypodermal IIS Transcriptional Targets

To investigate the molecular mechanisms of non-autonomous memory regulation from the hypodermis, we identified the downstream transcriptional targets regulated by hypodermal IIS in memory regulation. We compared the transcription profiles of Day 2 control (no auxin) and hypodermal DAF-2-degraded (+ auxin) worms. Principal component analysis (PCA) suggested a good separation of the two groups (**Figure 2A**). Consistent with our behavioral results, most genes upregulated by hypodermal IIS reduction are enriched in the nervous system and neurons, followed by the intestine and hypodermis (**Figure 2B**). Overall, 221 genes were significantly upregulated and 247 genes were significantly downregulated after DAF-2 was degraded in the hypodermis (log2 fold change ≥ 0.5 or ≤ -0.5, false discovery change [FDR] ≤ 1%) (**Figure 2C**, Supplementary Table S1).

**Figure 2.**
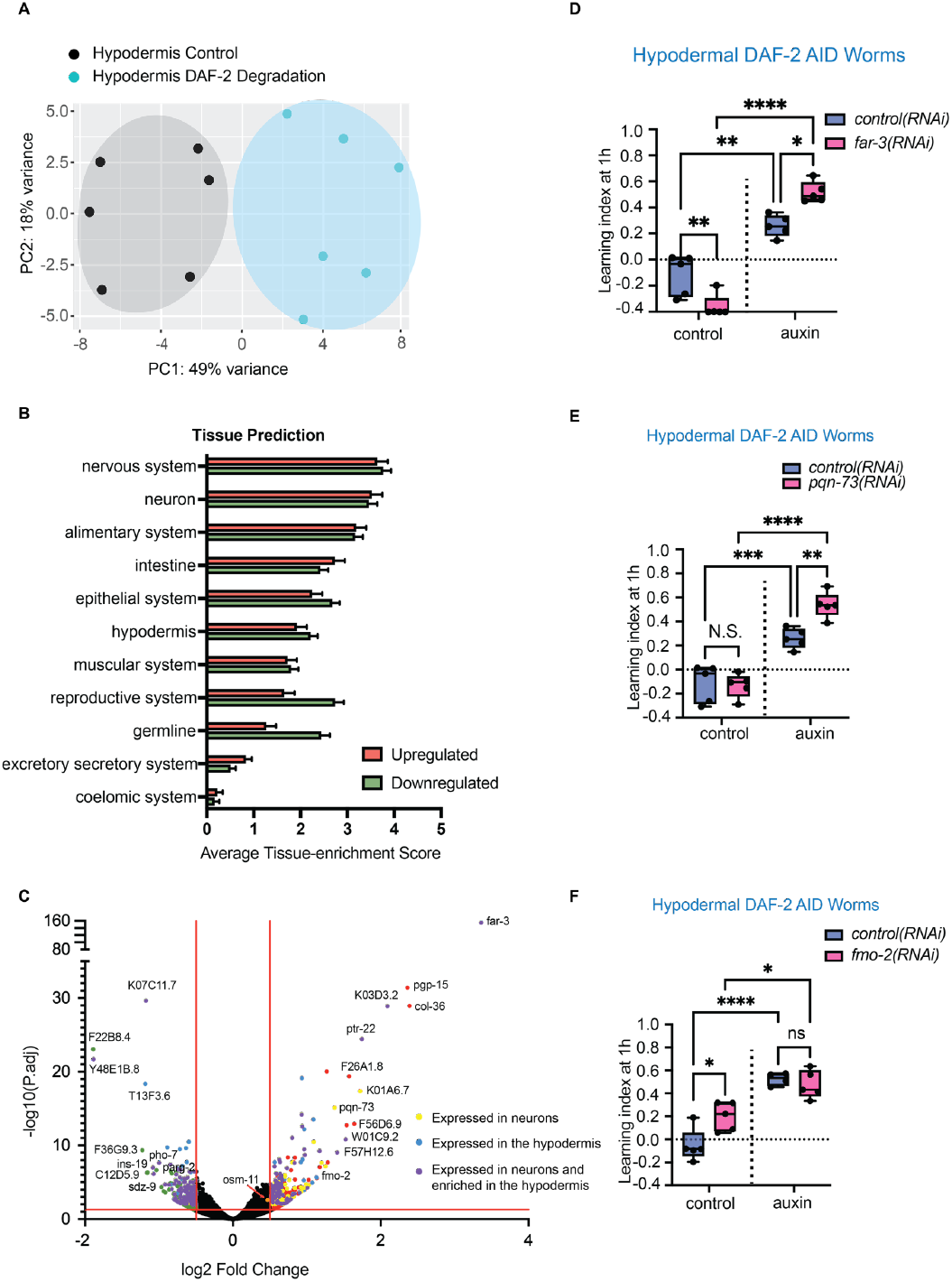
RNA-seq Transcriptional Profile of Whole Worms with Hypodermal DAF-2 Degradation Reveals Target Genes. (A) Principal component analysis (PCA) of the transcripts collected from Day 2 whole worms with hypodermal DAF-2 degradation (MQD2378, Day 2). Black dots: MQD2378 worms with no auxin treatment; Blue dots: MQD2378 worms with auxin treatment. (n=6 biological replicates per condition). (B) Prediction mean tissue enrichment scores for the genes significantly altered after hypodermal DAF-2 degradation. (C) Volcano plot of hypodermal DAF-2 (-) up- (red) and downregulated (green) genes (log2 fold change ≥ 0.5 or ≤ -0.5, false discovery change [FDR] ≤ 1%, n=6 biological replicates per condition). Yellow dots: genes that are expressed in neurons. Blue dots: genes that are expressed in the hypodermis. Purple dots: genes that are expressed in neurons and the hypodermis. (D-F) Knockdown of the upregulated genes, *far-3, pqn-73*, or *fmo-2*, did not block improved memory induced by hypodermal DAF-2 degradation (MQD2378, Day 3). n ≥ 4. (n represents the number of chemotaxis plates at each time point. Each plate contains ∼100 worms.) Two-way repeated measures ANOVA, Tukey post hoc tests. *p ≤ 0.05, **p ≤ 0.01, ***p ≤ 0.001, ****p ≤ 0.0001.

To test the target genes regulating memory downstream of hypodermal IIS, we examined whether knockdown of the top upregulated genes abrogated memory extension. *far-3*, the gene with the highest fold change, contains two DAF-16 binding elements (DBE) and one DAF-16 associative element (DAE) in its upstream promoter (420 bp, 408 bp and 978 bp). *far-3* encodes a fatty acid binding protein that is expressed in the hypodermis, intestine and neurons, and is regulated by Notch signaling key components^15,16^. *pqn-73* encodes a prion-like protein, and we previously found that *pqn-73* is expressed in arcade cells and neurons, and is required for long-term associative memory^5^. *fmo-2* encodes a flavin-containing monooxygenase expressed in the hypodermis and neurons, and activation of *fmo-2* in *C. elegans* extends lifespan^17^. However, *pqn-73* and *fmo-2* have no DBE or DAE in their promoters and are thus less likely to be direct targets of the insulin signaling pathway in the hypodermis (Supplementary Table S2). We found that knockdown of *far-3, pqn-73*, or *fmo-2* did not block memory enhancement induced by hypodermal DAF-2 degradation; in fact, their knockdown improved memory (**Figure 2D-F**), suggesting these genes were not responsible for the improved memory induced by hypodermal IIS reduction.

### The Notch Ligand OSM-11 is Required for the Non-autonomous Regulation of Memory

We next hypothesized that the direct hypodermal IIS targets might be diffusible proteins whose secretion is induced by hypodermal IIS reduction and then act non-autonomously in neurons and, from there, regulate memory. To identify diffusible candidate targets of the hypodermal IIS, we cross-referenced the database generated by Kaletsky et al., 2018, which revealed the tissue-specific gene expression in adult *C. elegans*^12^, with IIS Class I genes that were up-regulated in *daf-2* mutants compared to *daf-16;daf-2* worms^18,19^. We found 15 Class I candidate genes that are expressed in the hypodermis and are predicted to encode secreted proteins according to their amino acid sequences (**Figure 3A, B**). Among those 15 genes, *osm-11* had the highest hypodermal expression score^12^ (**Figure 3C**) and the highest fold of induction following adult hypodermal DAF-2 degradation (**Figure 3D**) as measured by RNAseq. Finally, *osm-11* has a DAF-16 binding Element (DBE) at 1078 bp and a DAF-16 associative Element (DAE) at 908 bp in its upstream promoter, indicating that *osm-11* may be a direct target of the hypodermal IIS.

**Figure 3.**
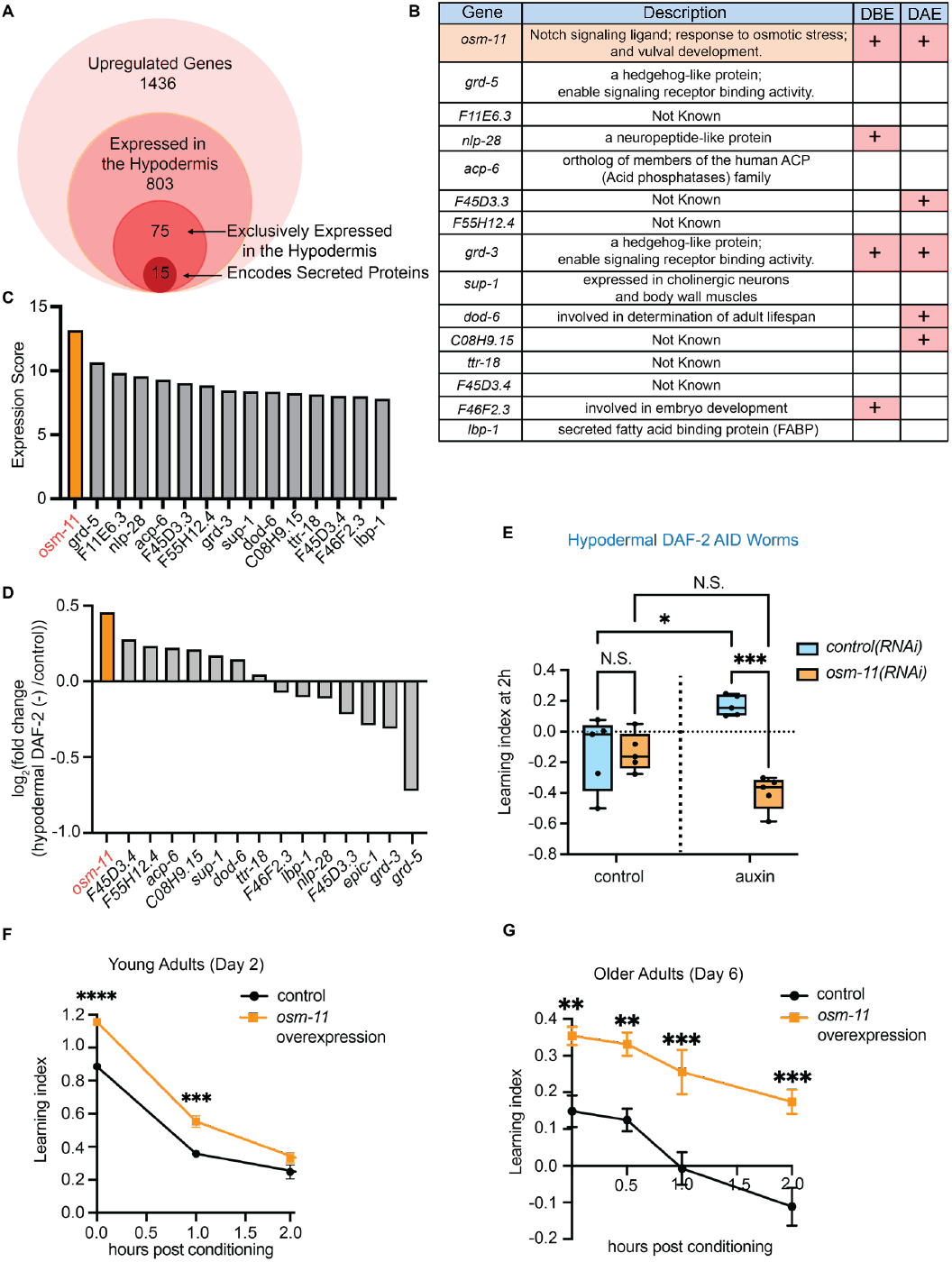
The Notch Ligand OSM-11 is Required for Memory Improvement Induced by Hypodermal DAF-2 Degradation. (A) 75 of 1436 Class I upregulated genes were exclusively expressed in the hypodermis. 15 genes are predicted to encode secreted proteins. (B) Description of all 15 candidate IIS targets. +: genes have DAF-16 Binding Element (DBE) or DAF-16 Associative Element (DAE) in their 1kb upstream promoters. (C) *osm-11* has the highest expression score in the hypodermis compared to the other 14 candidate targets. (D) log2 fold change shows the transcription level of the 15 candidate genes altered after hypodermal DAF-2 degradation. (E) Knocking down *osm-11*, which encodes a notch signaling ligand, by RNAi from L4 stage abolished the memory enhancement induced by hypodermal DAF-2 degradation in Day 3 adult worms (MQD2378). Representative experiment for 3 biological replicates, two-way repeated measures ANOVA, Tukey post hoc tests. (F) Overexpression of *osm-11* in young Day 2 adults (HA1133) enhanced memory. Representative experiment for 3 biological replicates. Mean ± s.e.m., two-way repeated measures ANOVA, Bonferroni post hoc tests. (G) Day 5 adults (HA1133) were heat-shocked for 1.5 hr at 33 degrees to induce the overexpression of *osm-11* and were tested for short-term associative memory after 24 hours of recovery. Overexpression of *osm-11* in aged Day 5 adults increased the memory of Day 6 worms. Representative experiment for 3 biological replicates. Mean ± s.e.m., two-way repeated measures ANOVA, Bonferroni post hoc tests. In all memory assays, n ≥ 4. (n represents the number of chemotaxis plates at each time point. Each plate contains ∼100 worms.) *p ≤ 0.05, **p ≤ 0.01, ***p ≤ 0.001, ****p ≤ 0.0001.

*osm-11* encodes a Notch ligand and is expressed in hypodermal seam cells and spermatheca of adult worms^20^. Therefore, we next asked whether the Notch ligand OSM-11 was required for the memory improvement we observed in animals with hypodermally-degraded DAF-2. Indeed, *osm-11* knockdown by RNAi abrogated the memory improvement induced by hypodermal DAF-2 degradation in young adults (**Figure 3E**), suggesting that reduced hypodermal IIS acts via the secreted protein OSM-11 to regulate short-term associative memory.

To determine if we can improve memory in wild-type worms by increasing Notch signaling, we used a heat-shock inducible strain to overexpress OSM-11 in young Day 2 adults and tested their short-term memory. OSM-11 overexpression increased short-term associative memory in wild-type worms (**Figure 3F**), while heat shock treatment did not affect short-term memory in control wild-type worms (Supplementary Figure S2). Furthermore, we found that heat shock induction of *osm-11* expression on Day 5 significantly enhanced memory in old Day 6 wild-type worms (**Figure 3G**) - an age when worms usually have no functional learning or memory (see control). These results suggest that not only is OSM-11 required for hypodermal-IIS memory enhancement, but also that OSM-11 overexpression is sufficient to enhance memory in the absence of additional inputs.

### The Notch Pathway Regulates Memory Improvement Induced by Hypodermal IIS Reduction

In *C. elegans*, Notch signaling is activated when the ligands bind to the receptors LIN-12 or GLP-1^21,22^, which are then cleaved, releasing the intracellular signaling transducing fragment, the Notch intracellular domain (NICD)^23,24^. NICD then translocates to the nucleus and binds to the transcription factor LAG-1 and coactivator LAG-3/SEL-8, forming a transcriptionally active ternary complex to regulate gene transcription^25^. Therefore, we hypothesized that OSM-11 might signal non-autonomously to activate neuronal Notch signaling to improve memory. If so, then other key components of Notch signaling should be required in neurons for the memory induced by hypodermal IIS reduction. We assessed short-term memory in hypodermal DAF-2 AID worms in which the Notch signaling components have been knocked down using RNAi. The GLP-1 Notch receptor is expressed in many neurons, including multiple ciliated sensory neurons^26^, but *glp-1* knockdown in ciliated neurons caused learning and chemotaxis defects (**Figure 4A**, Supplementary Figure S3A, B), preventing the analysis of subsequent short-term memory effects. The other Notch receptor, LIN-12, has been shown to be present in the RIG interneurons^27^. We tested *lin-12* knockdown in a subset of neurons using *flp-18p::lin12(RNAi)*, which drives the expression of a hairpin RNA in RIG, AIY, RIM and AVA interneurons^26^; knockdown of *lin-12* in those interneurons blocked memory enhancement induced by hypodermal IIS reduction (**Figure 4B**). Furthermore, knocking down the gene encoding the Notch signaling transcription factor LAG-1 in young adults blocked memory improvement upon hypodermal DAF-2 degradation (**Figure 4C**). Similarly, lack of the coactivator, LAG-3/SEL-8, in young hypodermal DAF-2-degraded adults prevented memory enhancement (**Figure 4D**), suggesting that multiple components of the Notch signaling pathway, including LIN-12, LAG-1, and SEL-8, are required for the extended memory induced by hypodermal IIS.

**Figure 4.**
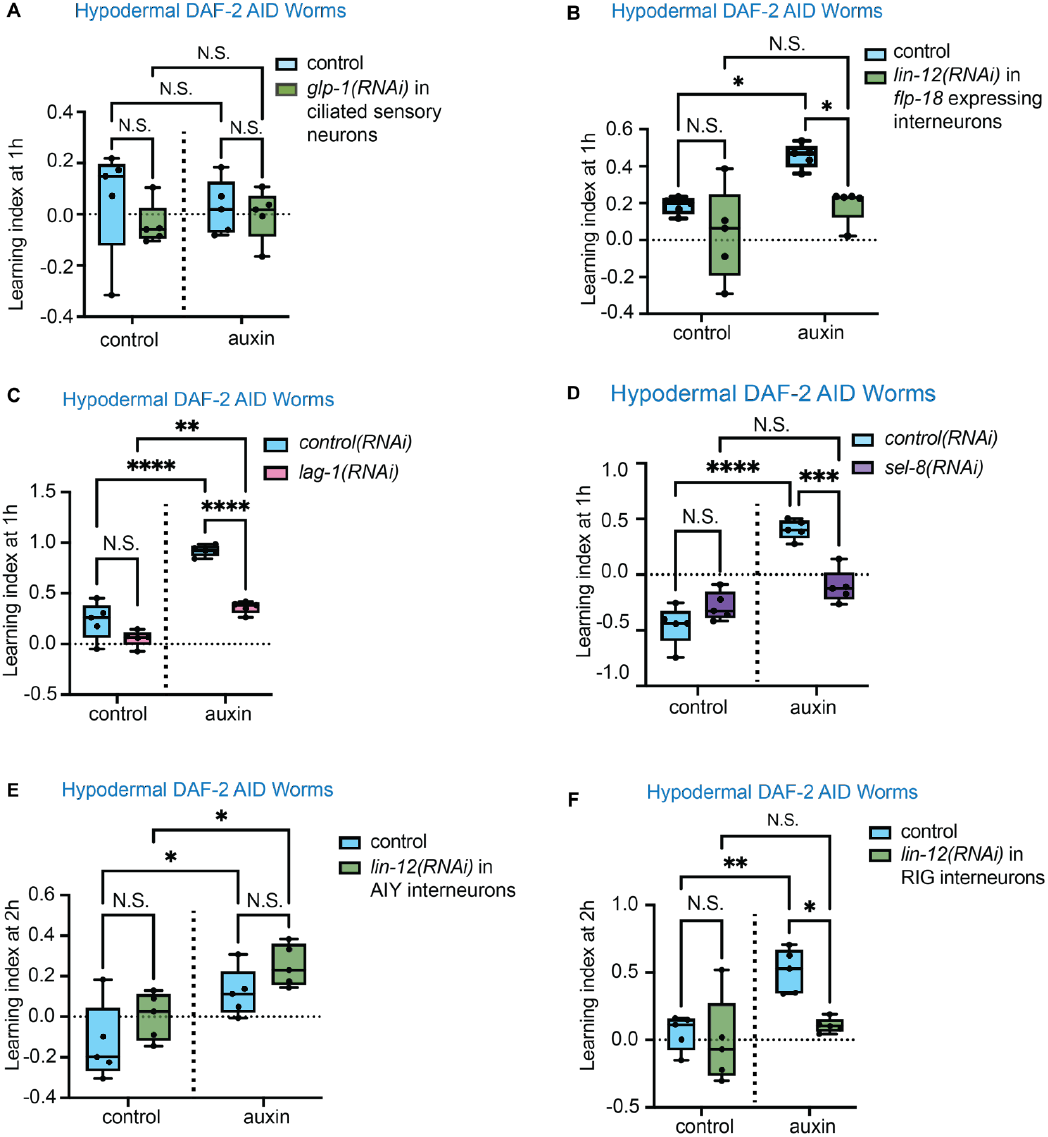
Notch Signaling in RIG interneurons is Required for Memory Improvement Enhanced by Hypodermal IIS Reduction. (A) Knocking down *glp-1*, one of the Notch signaling receptors, specifically in the ciliated neurons, abolished the ability to form short-term associative memory in Day 2 adult worms (CQ685). (B) Knocking down *lin-12*, one of the two Notch signaling receptors, specifically in *flp-18* expressed neurons (RIG, AIY, RIM, and AVA interneurons), abolished the memory enhancement induced by hypodermal DAF-2 degradation in Day 2 adult worms (CQ683). Representative experiment for 2 biological replicates. (C-D) Knocking down the Notch signaling transcription factor *lag-1* or the co-activator *sel-8*, by RNAi from L4 stage blocked the memory enhancement induced by hypodermal DAF-2 degradation in Day 3 adult worms (MQD2378). Representative experiment for 3 biological replicates. (E) Knocking down *lin-12*, specifically in the AIY interneurons, did not affect the memory enhancement induced by hypodermal DAF-2 degradation in Day 2 adult worms (CQ723). (F) Knocking down *lin-12*, specifically in the RIG interneurons, abolished the memory enhancement induced by hypodermal DAF-2 degradation in Day 2 adult worms (CQ721). Representative experiment for 3 biological replicates. All memory assay in Figure 4A-F used two-way repeated measures ANOVA, Tukey post hoc tests. n ≥ 4. (n represents the number of chemotaxis plates at each time point. Each plate contains ∼100 worms.) *p ≤ 0.05, **p ≤ 0.01, ***p ≤ 0.001, ****p ≤ 0.0001.

To determine whether the Notch pathway is also required for neuron-autonomous IIS-regulated memory, we tested the effect of *sel-8(RNAi)* on the neuronal DAF-2 AID worms’ memory. Knocking down *sel-8* had no effect on memory improvement induced by neuronal-DAF-2 degradation (Supplementary Figure S3C), suggesting that reduced neuronal DAF-2 increases short-term memory through distinct mechanisms that may not involve Notch signaling.

To further identify in which interneurons Notch signaling acts, we tested hypodermal IIS degron worms with *lin-12* knocked down in subsets of the interneurons affected by the *flp-18* promoter, including RIM, AVA, AIY, and RIG^27–30^. *nmr-1p::lin-12(RNAi)* worms were not healthy, preventing analysis of the effect of *lin-12* knockdown in RIM and AVA in memory. The knockdown of *lin-12* in AIY neurons (*ttx-3p::lin- 12(RNAi)*), an interneuron that is required for long-term associative memory formation in wild-type worms^31^, did not abrogate memory improvement (**Figure 4E**). *lin-12* knockdown in RIG interneurons (*twk-3p::lin-12(RNAi)*) significantly blocked memory enhancement by hypodermal IIS reduction (**Figure 4F**), suggesting that the Notch signaling in RIG interneurons is downstream of OSM-11. Taken together, these results suggest that hypodermal IIS acts through Notch signaling via RIG interneurons in short-term memory regulation, and uses a mechanism that is separate from that of neuronal insulin/IGF-1 signaling regulation of memory.

### The Notch Target, *ins-19*, Regulates Memory Extension in a DAF-16-dependent Manner

The function of Notch signaling is to regulate gene expression. In our transcriptional analysis of genes regulated upon degradation of DAF-2 in the hypodermis, we found that many of the targets (24 up and 38 down) have Notch binding sites in their promoters, and 36 of these genes are predicted to be neuronally expressed (Supplementary Table S2). The insulin-like peptide gene *ins-19* was significantly downregulated upon hypodermal DAF-2 degradation; its promoter does not have a DAF-16 binding element (DBE) or DAF-16 associate element (DAE), indicating that *ins-19* is most likely an indirect target of hypodermal IIS. In fact, the *ins-19* promoter has two Notch binding sites (Supplementary Figure S3D). Expression of *ins-19* is low in young adults, but increases with age (Weng and Murphy, unpublished; **Figure 5A**). The fact that *ins-19* is downregulated upon hypodermal DAF-2 degradation suggested its reduction might be sufficient to extend STAM. Indeed, we found that *ins-19(tm5155)* mutants exhibited enhanced short-term memory (**Figure 5B**). We mutated the two Notch binding sites and found that homozygous mutants are embryonic lethal, suggesting that *ins-19* expression is critical for development; therefore, we were unable to test this mutant in adulthood for its effect on STAM. However, mutation of *daf-16* in the *ins-19* background (*daf-16(mu86);ins-19(tm5155)*) at least partially impaired STAM (**Figure 5C**), suggesting that the INS-19 peptide is likely a neuronally-expressed insulin-like agonist for DAF-2^32^, engaging the neuronal DAF-2 STAM pathway (**Figure 1A**). Suppression of this agonist might help slow the normal increase in DAF-2 activation with age and thus loss of memory with age. Thus, Notch signaling from the hypodermis downstream of DAF-2/DAF-16 signaling regulates STAM through its control both of direct Notch-regulated targets and through insulin/DAF-2/DAF-16 signaling in neurons (**Figure 5D**). Taken together, these results suggest that hypodermal IIS acts through Notch signaling via RIG interneurons in short-term memory regulation.

**Figure 5:**
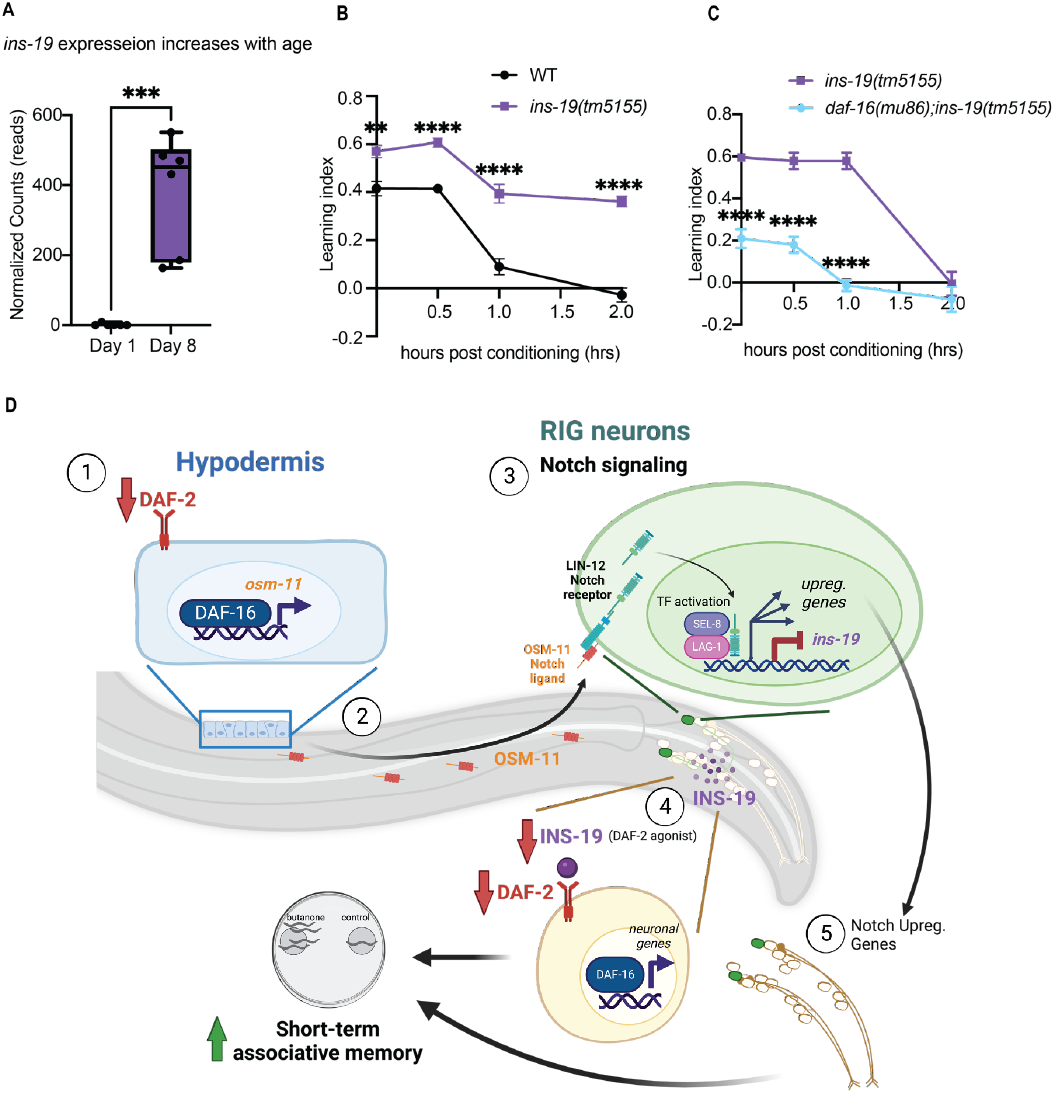
One of the Notch Targets, *ins-19*, Regulates Memory Extension in a DAF-16-dependent Manner and the Model of Hypodermal IIS – Notch Regulation of Memory. (A) *ins-19* transcription levels increased with age in wild-type neurons. Gene transcription changes with age were measured using RNAseq on isolated neurons from Day 1 and 8 wild-type worms. (n=6 biological replicates). (B) Day 3 *ins-19* mutants (QL188) exhibited increased STAM compared to wild-type worms. Representative experiment for 3 biological replicates. (C) *daf-16* mutation at least partially suppressed this increased STAM in *ins-19(tm5155)* background. For the memory assays in 5B-C, Mean ± s.e.m., n ≥ 4. (n represents the number of chemotaxis plates at each time point. Each plate contains ∼100 worms.) **p ≤ 0.01, ****p ≤ 0.0001, two-way repeated measures ANOVA, Bonferroni post hoc tests. (D) (1) The reduction of hypodermal IIS promotes the transcription factor FOXO/DAF-16 to translocate into the nucleus and upregulate *osm-11* gene expression. 2) OSM-11 then diffuses to the RIG interneurons and 3) activates the neuronal Notch signaling by binding to the Notch receptor LIN-12 and GLP-1. The activated Notch receptor is cleaved, and the intracellular domain transfers into the nucleus and forms a complex with the transcription factor LAG*-*1 and the coactivator SEL-8 to regulate many downstream Notch target gene transcription. 4) Among those genes, the downregulation of *ins-19*, an IIS agonist, may function to suppress DAF-2 in other neurons, and 5) together with other Notch signaling regulated neuronal genes, extend memory.

## Discussion

In this study, we present evidence of non-autonomous memory regulation mediated by the hypodermal insulin pathway. Previously it was found that *C. elegans* with elevated insulin signaling have learning impairments associated with high neuronal levels of kynurenic acid, whose precursor is generated in epidermal cells^33,34^, indicating some involvement of non-neuronal tissues in memory regulation; however, whether reduced insulin signaling in non-neuronal tissues could benefit learning and memory was not yet known. We first found that DAF-2 degradation exclusively in the hypodermis improves short-term memory, and our transcriptomic results and memory assay results suggest that the upregulation of a diffusible Notch ligand, OSM-11, mediates this memory improvement. We further identified key downstream components of the Notch signaling pathway, including the Notch receptor LIN-12 in RIG neurons, the transcription factor LAG-1, and the co-activator SEL-8, that are required for memory improvement induced by hypodermal DAF-2 degradation (**Figure 5D**). Furthermore, reducing the IIS pathway in the hypodermis or increasing Notch signaling in aged worms can each improve short-term memory. Together, the results suggest a novel mechanism for regulation of memory by a non-neuronal tissue, the hypodermis.

The improved cognitive function enabled by the hypodermal tissue presents a new angle for treatment to slow cognitive aging. Notch signaling is an evolutionarily conserved pathway, and its roles in development, cancer, and nervous system development^35,36^, as well as its role in *C. elegans* development, have been studied extensively^37–39^. However, Notch signaling’s regulation of memory in *C. elegans* was previously unknown. Our finding that neuronal Notch signaling regulates memory in both young and aged adult *C. elegans* is consistent with findings in flies and mice: the post-developmental inactivation of Notch signaling in Drosophila causes deficits in long-term memory (LTM)^40^, and a transient increase of Notch signaling can facilitate LTM^41^. Postnatal disruption of Notch signaling in the mouse brain causes defects in learning and memory^42,43^. Together with previous findings from other invertebrates and mammals, Notch signaling appears to have a conserved function in memory regulation from *C. elegans* to mammals.

The *C. elegans* hypodermis serves not only to facilitate cuticle formation during development and to sense external environmental changes such as shortage of food and osmolarity^12,44,45^, but it is also essential for metabolic functions^12^. RIG neurons regulate CO^2^ response during the early stage of food deprivation^46^; the fact that LIN-12 is required in the RIG neurons for OSM-11-mediated memory improvement may indicate that the hypodermis might facilitate worms’ sensing of food and other environmental cues. The *C. elegans* hypodermis is one of the worm’s largest metabolic tissues, and shares transcriptional similarity with human liver and blood plasma^12^. Interestingly, there may be a parallel in the effect of the mammalian liver on memory: the administration of systemic blood plasma from exercised aged mice – which secrete factors from the liver into the blood - improves cognitive function and neurogenesis in sedentary aged recipient mice^47^. Likewise, Alzheimer’s disease model rats receiving administration of a potent metabolic regulator, fibroblast growth factor 21 (FGF21), mainly produced by the liver, also exhibit ameliorated neuro-degeneration and enhanced learning and memory^48^. It is possible that the worm’s hypodermis may function somewhat like the liver: the hypodermis’ non-autonomous regulation of memory might allow worms to respond more readily to future environmental changes, with different temporal dynamics than regulation in the neurons. By engaging insulin signaling in the neurons downstream of Notch signaling, the worms may be able to have a more sustained memory response than insulin signaling alone might achieve. Together, these data reveal a critical connection between metabolism and memory regulation, suggesting a similar mechanism across different animal models and providing new therapeutic insight to ameliorate cognitive aging or to treat neurodegenerative diseases by regulating metabolism.

## Acknowledgements

We thank the *Caenorhabditis* Genetics Center (CGC) and the National BioResource Project (NBRP) for strains, Jasmine Ashraf, William Keyes, and Morgan Stevenson for help with experiments, members of the Murphy Lab for discussion and feedback on the manuscript, Biorender.com for graphical abstract and model figure design software. CTM is the Director of the Simons Collaboration on Plasticity in the Aging Brain (SCPAB), which supported the work, and the Glenn Center for Aging Research at Princeton. SZ was supported by China Scholarship Council (CSC). This project was funded in part by the National Natural Science Foundation of China (NSFC-ISF 32061143020 to M-QD).

## Methods

### Bacteria Strains

| Strain | Source |
| --- | --- |
| *E. coli*: OP50 | BactoBeads™ |
| *E. coli*: HT115 | Caenorhabditis Genetics Center |

### *C. elegans* strains

#### General worm maintenance

***C. elegans* strains****General worm maintenance**
| Strain | Source | Identifier | Ref |
| --- | --- | --- | --- |
| N2 var. Bristol: wild type | Caenorhabditis Genetics Center | Wild-type |  |
| *frSi21 [col-62p::rde-1 3'UTR] II. frIs7 [nlp-29p::GFP + col-12p::DsRed] IV* | Caenorhabditis Genetics Center | IG1846 |  |
| *unc-119(ed3); hqKi373 [Prgef-1::TIR1::mRuby::unc-54 3'UTR, cb-unc-119(+)] II; daf-2(hqKi363[daf-2::degron::mNeonGreen]) III* | Dong lab | MQD2356 | 14 |
| *unc-119(ed3); ieSi61 [Pges-1::TIR1::mRuby::unc-54 3'UTR, cb-unc-119(+)] II; daf-2(hqKi363[daf-2::degron::mNeonGreen]) III* | Dong lab | MQD2374 | 14 |
| *unc-119(ed3); hqKi378 [Plim-7::TIR1::mRuby::unc-54 3'UTR, cb-unc-119(+)] II; daf-2(hqKi363[daf-2::degron::mNeonGreen]) III* | Dong lab | MQD2383 | 14 |
| *daf-2(hqKi363[daf-2::degron::mNeonGreen]) III; ieSi38 [Psun-1::TIR1::mRuby::sun-1 3'UTR, cb-unc-119(+)] IV* | Dong lab | MQD2375 | 14 |
| *unc-119(ed3); hqKi374 [Pdpy-7::TIR1::mRuby::unc-54 3'UTR, cb-unc-119(+)] II; daf-2(hqKi363[daf-2::degron::mNeonGreen]) III* | Dong lab | MQD2378 | 14 |
| *unc-119(ed3); hqKi375 [Pmyo-3::TIR1::mRuby::unc-54 3'UTR, cb-unc-119(+)] II; daf-2(hqKi363[daf-2::degron::mNeonGreen]) III* | Dong lab | MQD2379 | 14 |
| *rtIs26[hsp::osm-11, elt-2p::gfp, pha-1(+)]III* | Hart lab | HA1133 | 26 |
| *rtEx721[flp-18p::lin-12(RNAi) myo-2p::gfp]; glp-1(bn18)III* | Hart lab | HA2160 | 26 |
| *rtEx635[che-2p::glp-1(RNAi), myo-2p::gfp pha-1(+)]; pha(e2123)III* | Hart lab | HA1931 | 26 |
| *daf-2(e1370) III* | Shen's lab | CB1370 |  |
| *wqEx76 [Ptwk-3::sense:lin-12, Ptwk-3::antisense:lin-12 + Pmyo-2::GFP]* | This paper | CQ721 |  |
| *wqEx77 [Pttx-3::sense:lin-12, Pttx-3::antisense:lin-12 + Pmyo-2::GFP]* | This paper | CQ723 |  |
| *wqEx78 [Pnmr-1::sense::lin-12, Pnmr-1::antisense::lin-12 + Pmyo-2::GFP]* | This paper | CQ734 |  |
| *unc-119(ed3); hqKi374 [Pdpy-7::TIR1::mRuby::unc-54 3'UTR, cb-unc-119(+)] II; daf-2(hqKi363[daf-2::degron::mNeonGreen]) III; rtEx721[flp-18p::lin-12(RNAi) myo-2p::gfp]; glp-1(bn18)III* | This paper | CQ683 |  |
| *unc-119(ed3); hqKi374 [Pdpy-7::TIR1::mRuby::unc-54 3'UTR, cb-unc-119(+)] II; daf-2(hqKi363[daf-2::degron::mNeonGreen]) III; rtEx635[che-2p::glp-1(RNAi), myo-2p::gfp pha-1(+)]; pha(e2123)III* | This paper | CQ685 |  |
| *ins-19(tm5155); daf-16(mu86)* | This paper | CQ726 |  |

All strains were cultured using standard methods^49^. For all experiments, worms were maintained at 20°C on plates made from nematode growth medium (NGM: 3 g/L NaCl, 2.5 g/L Bacto-peptone, 17 g/L Bacto-agar in distilled water, with 1 mL/L cholesterol (5 mg/mL in ethanol), 1 mL/L 1M CaCl2, 1 mL/L 1M MgSO4, and 25 mL/L 1M potassium phosphate buffer (pH 6.0) added to molten agar after autoclaving) or high growth medium (HGM: NGM recipe modified as follows: 20 g/L Bacto-peptone, 30 g/L Bacto-agar, and 4 mL/L cholesterol (5 mg/mL in ethanol); all other components same as NGM), with OP50 *E. coli* for *ad libitum* feeding. To synchronize experimental animals, eggs were collected from gravid hermaphrodites by exposing the animals to an alkaline-bleach solution (e.g., 1.5 ml sodium hypochlorite, 0.5 mL 5 N KOH, 8.0 mL water), followed by repeated washing of collected eggs in M9 buffer ((6 g/L Na2HPO4, 3 g/L KH2PO4, 5 g/L NaCl and 1 mL/L 1M MgSO4 in distilled water).

#### Sequences of the primers used for constructing transgenes that knockdown *lin-12* in AIY, RIG, or RIM and AVA interneurons

Three promoters were used to drive the sense and antisense expression of *lin-12* RNA: p*ttx-3* (AIY interneurons), p*twk-3* (RIG interneurons), and p*nmr-1* (RIM and AVA interneurons).

For the *ttx-3* promoter, a 3.1 kb fragment upstream of the ATG and the first two exons and introns were amplified using the following primers:

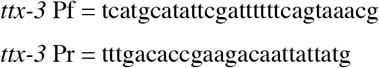

For the *twk-3* promoter, a 149 bp fragment upstream of the ATG was amplified using the following primers:

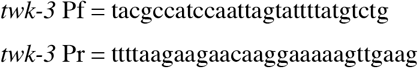

For the *nmr-1* promoter, a 5 kb fragment upstream of the ATG was amplified using the following primers:

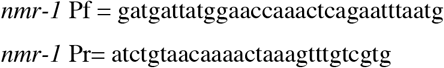

Because these promoters were used to drive transcription of *lin-12* gene, all three Pr of primers had 25 additional nucleotides complementary to either the sensed strand or the antisense strand of *lin-12* gene.

3.8 kb of the target gene *lin-12* was amplified from the strain HA2160 using the following primers:

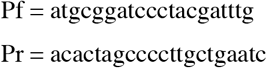

The fragments for *twk-3p*- or *ttx-3p*-driven the sense and antisense expression of *lin-12* RNA were injected at 100 ng/ul each with 2 ng/ul myo-2p ::GFP marker at a total volume of 20 ul. The fragments for *nmr-1p*-driven the sense and antisense expression of *lin-12* RNA were injected at 75 ng/ul each with 2 ng/ul myo-2p ::GFP marker and 1 kb DNA ladder to bring up to 100 ng/ul at a total volume of 20 ul.

#### Auxin treatment

For auxin experiments, the standard HGM molten agar was supplemented with 2.5 mL 400 mM indole-3-acetic acid (IAA): Alfa Aesar (#A10556) freshly prepared in ethanol and plates were seeded with *E. coli* for *ad libitum* feeding. Synchronized AID worms were transferred to HGM with auxin plates for 16-20 hours before behavior assay. For mid-life experiments, worms were transferred at the L4 larval stage onto HGM plates supplemented with 500 ml/L 0.1 M FUDR (5-Fluoro2’-deoxyuridine) for a final concentration of 0.05M FUDR and were transferred back to standard HGM with or without auxin 20 hours before memory assay.

#### RNAi treatment

For RNAi experiments, the standard HGM molten agar was supplemented with 1 mL/L 1 M IPTG (isopropyl β-d-1-thiogalactopyranoside) and 1 mL/L 100 mg/mL carbenicillin, and plates were seeded with HT115 *E. coli* for ad libitum feeding. All RNAi treatment starts from L4 larval stage and lasts for 2-3 days. RNAi experiments were performed using the standard feeding RNAi method. Bacterial clones expressing the control construct (empty vector, pL4440). All RNAi clones were sequenced prior to use.

| RNAi | Source |
| --- | --- |
| *L4440* | Addgene (Fire Lab) |
| *pAD-12* | Addgene (Kenyon Lab) |
| *daf-2* | Ahringer |
| *far-3* | Ahringer |
| *pqn-73* | Ahringer |
| *fmo-2* | Horizon Discovery |
| *osm-11* | Ahringer |
| *lag-1* | Horizon Discovery |
| *sel-8* | Ahringer |

#### Short-Term (Pavlovian) Associative Assay (STAM)

Animals were trained and tested for short-term memory as previously described^3^. Briefly, synchronized young or aged adult hermaphrodites were washed from HGM, RNAi or HGM supplemented with auxin plates with M9 buffer, allowed to settle by gravity, and repeatedly washed three more times with M9 buffer. Then animals were starved for 1 hr in M9 buffer. For conditioning (food and 10% 2-butanone pairing), worms were then transferred to 10 cm NGM conditioning plates (seeded with OP50 *E. coli* bacteria and with 18 mL 10% 2-butanone (Acros Organics) dissolved in ethanol on the lid) for 1 hr at 20°C. After conditioning, the trained worms were tested for chemotaxis towards 10% butanone vs. an ethanol control either immediately (0 hr) or after being transferred to 10 cm NGM holding plates with fresh OP50 *E. coli* bacteria for specified time intervals (30 min-2 hr).

Chemotaxis indices were calculated as follows:

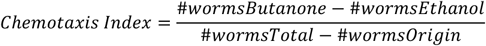

The calculation for the learning Index is:

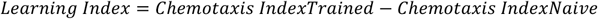

Learning indices for extrachromosomal transgenic strains were analyzed by hand counting GFP positive and negative worms at different locations at individual timepoint on the chemotaxis plates. Control animals for these experiments were the transgenic worms’ GFP-negative siblings.

#### Chemotaxis Assay

Synchronized adult worms were tested for chemotaxis to 1% benzaldehyde in ethanol or 10% pyrazine in ethanol, using standard, previously described chemotaxis assay conditions^50^.

#### Heat shock transgene induction

Transgenic worms were cultivated on standard HGM plates (or HGM with FUdR for mid-life experiments) seeded with OP50 bacteria until behavior assay (or transferred to standard HGM plates with OP50 24 hours before behavior assay for mid-life experiments). The plates were placed (agar-side down) in an incubator at 33°C for 90 min. For young animals, worms were recovered at least 2 hours at 20°C before behavior assay. For mid-life experiments, worms were recovered for 24 hours at 20°C before behavior assay.

#### Imaging

Worms were treated with 1 mM auxin for 24 hours starting from late L4 stage at 20 °C. Confocal images were captured by the spinning-disk microscope (UltraVIEW VOX; PerkinElmer) equipped with a 60×, 1.4 numerical aperture oil-immersion objective. Images were viewed and processed using Volocity software (PerkinElmer).

#### RNA isolation

Synchronized Day 2 adult worms were collected in M9 and washed repeatedly 3 times to remove excess bacteria. Worm pellets were crushed in liquid nitrogen and transferred to 850 uL Trizol LS. Total RNA was extracted using the standard Trizol/chloroform/isopropanol method followed by DNase digestion and Qiagen RNeasy Mini kit. Agilent Bioanalyzer RNA Pico chips were used to assess the quality and quantity of isolated RNA. mRNA libraries were prepared using the RNA-Seq directional library prep on Apollo 324 robot and were sequenced (65-nt paired-end) on the Illumina Novaseq S1 100nt flowcell v1.5 platform (yields ∼1.3-1.6 B reads).

#### RNA-seq data analysis

RNA sequencing analysis was performed as previously described. FASTQC was used to assess read quality scores. The universal Illumina adaptor sequences were trimmed using Cutadapt v1.6 (Martin, 2011). The trimmed reads were mapped to the *C. elegans* genome (UCSC Feb 2013, ce11/ws245) using STAR (Dobin et al., 2013). The reads aligning to individual genes were counted using htseq-counts (mode: union), and DESeq2 was used for differential expression analysis. Genes with an adjusted *p-value* ≤ 0.05 were considered significantly differentially expressed.

#### Tissue Query

worm.princeton.edu was used for tissue query from upregulated or downregulated gene lists (DESeq2 genes FDR < 1%).

#### Secreted Protein Prediction

SignalP-5.0 (DTU Health Tech) was used for secreted protein prediction from 75 genes that are exclusively expressed in the hypodermis.

#### Quantification and Statistical Analysis

Two-way ANOVA with Tukey post hoc tests was used to compare learning indices among multiple groups. one-way ANOVA analysis of variances followed by Bonferroni post hoc tests for multiple comparisons was performed. Two-way ANOVAs were used between genotype (*daf-2(e1370)* and wild-type, control RNAi and *daf-2* RNAi, control and *osm-11* overexpression, wild-type and *ins-19(tm5155), ins-19(tm5155 and daf-16(mu86);ins-19(tm5155))* and different timepoint (0 hr, 0.5 hr, 1 hr, 2 hr) on learning indices with a significant interaction between factors leading to the performance of Bonferroni post hoc comparisons to determine differences between individual groups. Experiments were repeated on separate days, using separate independent populations, to confirm that results were reproducible. Prism 9 software was used for all statistical analyses. Software and statistical details used for RNA sequencing analyses are described in the method details section of the STAR Methods. Additional statistical details of experiments, including sample size (with n representing the number of chemotaxis assays performed for behavior, RNA collections for RNA-seq, and the number of worms for microscopy), can be found in the figure legends.

## Supplementary Tables and Figures

**Supplementary Table S1. RNA-seq Results of Hypodermal DAF-2 (-) Worms**

Significantly up-and down-regulated genes from RNAseq analysis of Day 2 hypodermal DAF-2 AID worms with auxin vs without auxin treatment.

**Supplementary Table S2. Expression pattern and DBE, DAE, and Notch binding motifs of the differentially expressed genes**.

Tissue expression pattern of the differentially expressed genes according to Wormbase and Kaletsky. et al., 2018. DBE, DAE, and Notch binding motifs of the differentially expressed genes according to their sequences.

**Supplementary Figure S1.**
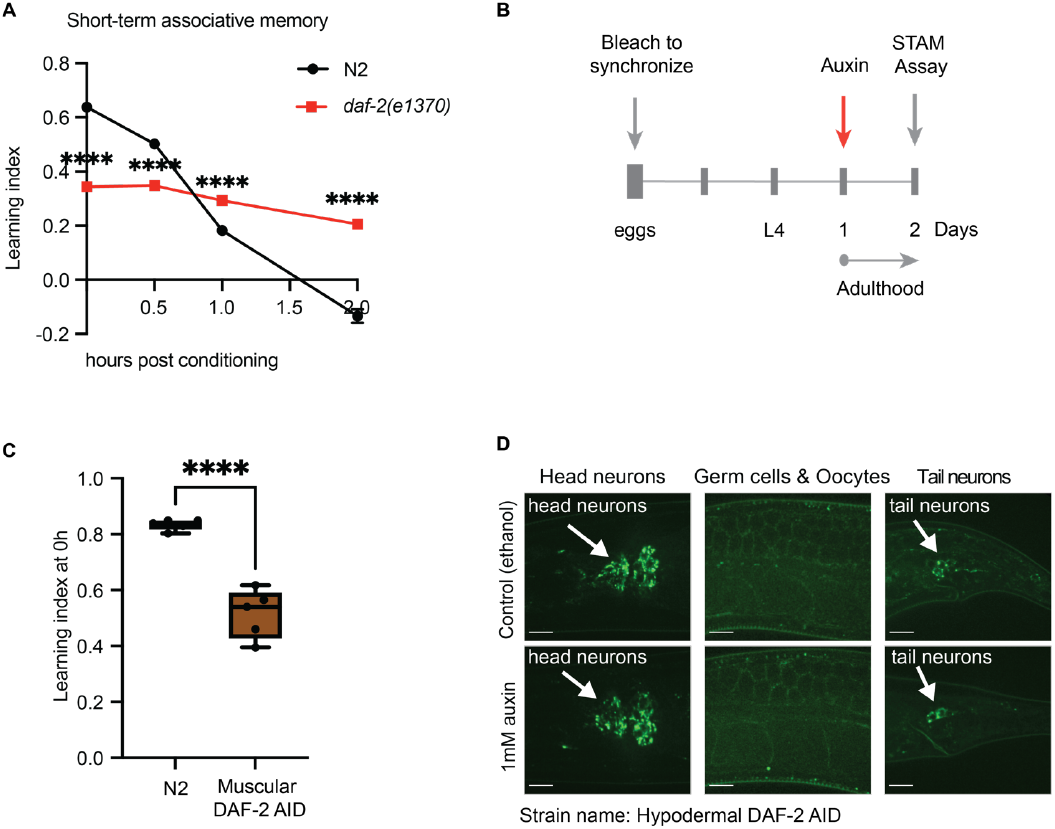
*daf-2* Mutants Had Improved Short-term Associative Memory and Knocking Down *daf-2* Specifically in the Hypodermis Improved Memory. (A) *daf-2* mutants had improved short-term associative memory. Mean ± s.e.m., n ≥ 4. (n represents the number of chemotaxis plates at each time point. Each plate contains ∼100 worms.) ****p ≤ 0.0001, two-way repeated measures ANOVA, Bonferroni post hoc tests. (B) Auxin treatment started from Day 1, and STAM assay was performed on Day 2. (C) Worms with auxin-inducible DAF-2 degradation in the muscle (MQD2379) had learning defects prior to auxin treatment in Day 2 adult animals. n ≥ 4. (n represent the number of chemotaxis plates at each time point. Each plate contains ∼100 worms.) ****p ≤ 0.0001. Unpaired t-test. (D) *Pdpy-7 ::TIR-1* did not induce DAF-2 degradation in neurons. DAF-2 is fused with a mNeonGreen tag at the C-termini. White arrows point to head and tail neurons. Scale bar: 10 μm.

**Supplementary Figure S2.**
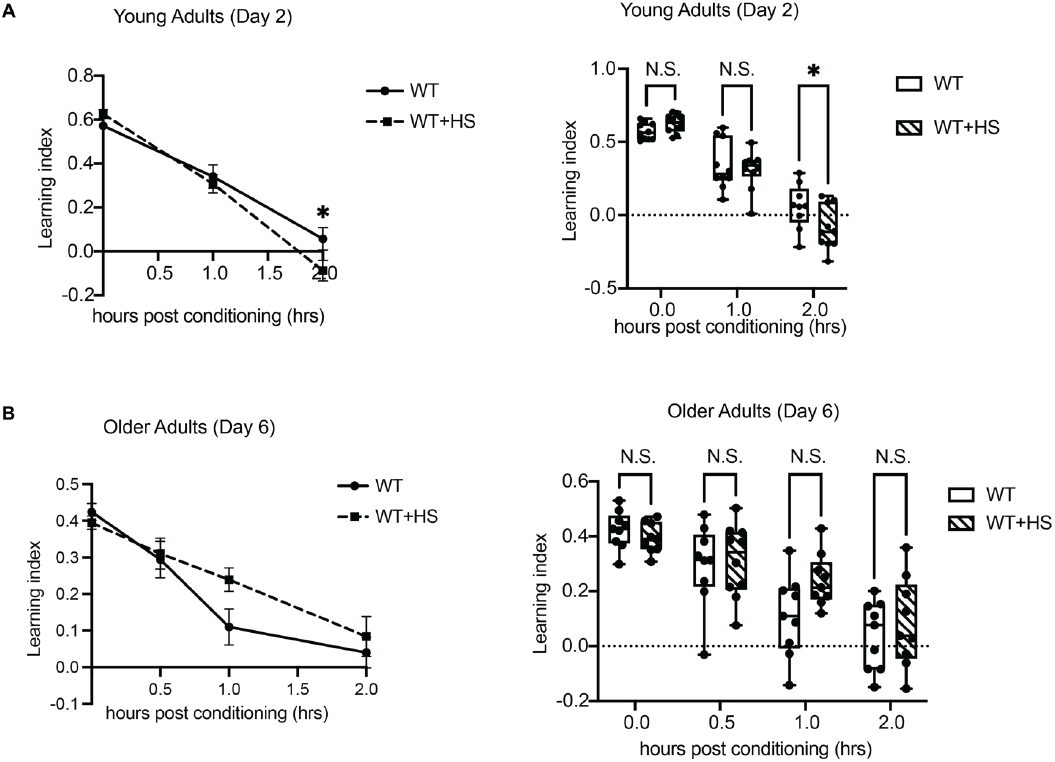
Heat Shock Had No Effect on Short-term Associative Memory in Both Young and Old Wild-type Worms. (A) Heat shock had no effect on short-term associative memory in young Day 2 wild-type worms. Day 2 young wild-type adults were heat-shocked for 1.5 hr at 33 degree and were tested for short-term associative memory by STAM assay after rest for 2 hours. (B) Heat shock had no effect on short-term associative memory in old Day 6 wild-type worms. Day 5 old wild-type adults were heat-shocked for 1.5 hr at 33 degree and were tested for short-term associative memory by STAM assay after rest for 24 hours. Mean ± s.e.m., n ≥ 9. (n represents the number of chemotaxis plates at each time point. Each plate contains ∼100 worms.) Two-way repeated measures ANOVA, Bonferroni post hoc tests.

**Supplementary Figure S3.**
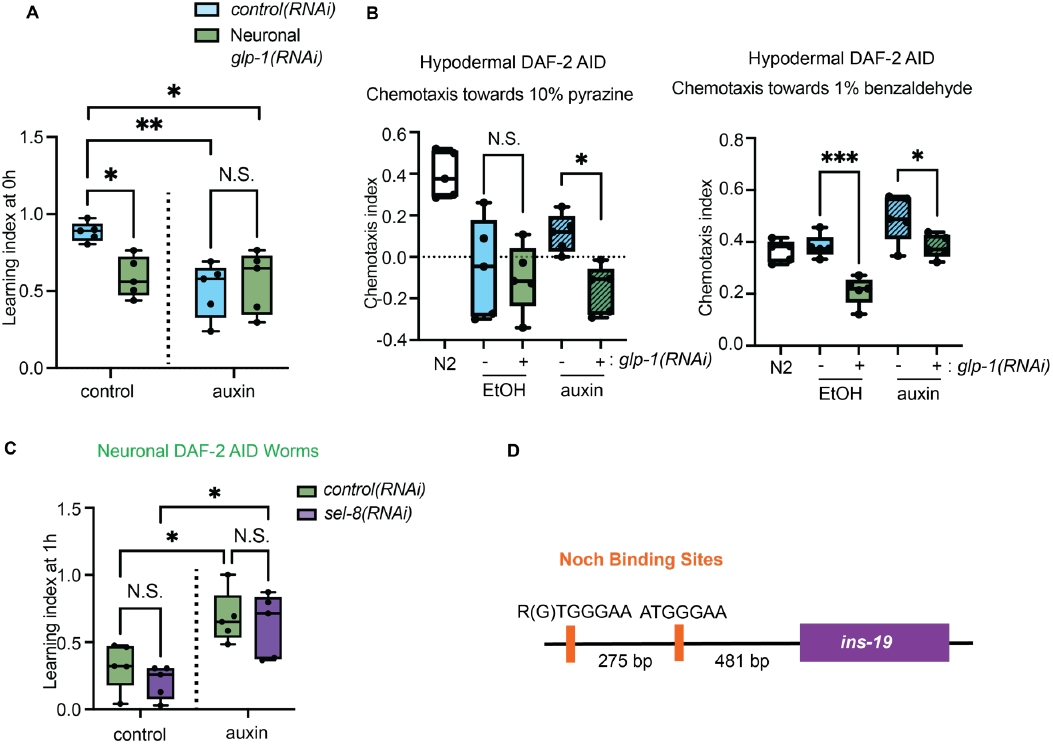
Knocking Down *glp-1* Specifically in Ciliated Neurons Impaired Chemosensory and Notch binding sites of *ins-19*. (A) Knocking down *glp-1*, one of the Notch signaling receptors, specifically in the ciliated neurons, abolished the ability to learn in Day 2 adult worms, thus STAM cannot be measured. n ≥ 4. (n represents the number of chemotaxis plates at each time point. Each plate contains ∼100 worms.) *p ≤ 0.05. **p ≤ 0.01, two-way repeated measures ANOVA, Tukey post hoc tests. (B) Knocking down *glp-1* in the ciliated neurons impaired chemotaxis towards 10% pyrazine and 1% benzaldehyde in young Day 2 adult worms. n ≥ 4. (n represents the number of chemotaxis plates at each time point. Each plate contains ∼100 worms.) *p ≤ 0.05, ***p ≤ 0.001. Unpaired t-test. (C) The Notch signaling was not required for memory enhanced by neuronal DAF-2 degradation. Knocking down *sel-8* by RNAi from L4 stage did not affect the memory enhanced by loss of neuronal DAF-2 (MQD2356). n ≥ 4. (n represents the number of chemotaxis plates at each time point. Each plate contains ∼100 worms.) *p ≤ 0.05, two-way repeated measures ANOVA, Tukey post hoc tests. (D) Two Notch binding sites are found (488 bp and 770 bp) upstream of *ins-19*.

